# Mutational analysis of field cancerization in bladder cancer

**DOI:** 10.1101/536466

**Authors:** Trine Strandgaard, Iver Nordentoft, Philippe Lamy, Emil Christensen, Mathilde Borg Houlberg Thomsen, Jørgen Bjerggaard Jensen, Lars Dyrskjøt

**Affiliations:** Department of Molecular Medicine, Aarhus University Hospital, 8200 Aarhus N, Denmark; Department of Clinical Medicine, Health, Aarhus University, 8000 Aarhus C, Denmark; Department of Urology, Aarhus University Hospital, 8200 Aarhus N, Denmark

## Abstract

The multifocal and recurrent nature of bladder cancer has been explained by field cancerization of the bladder urothelium. To shed light on field cancerization in the bladder, we investigated the mutational landscape of normal appearing urothelium and paired bladder tumors from four patients. Sequencing of 509 cancer driver genes revealed the presence of 2-16 mutations exclusively localized in normal tissue (average target read depth 634x). Furthermore, 6-13 mutations were shared between tumor and normal samples and 8-75 mutations were exclusively detected in tumor samples. More mutations were observed in normal samples from patients with multifocal disease compared to patients with unifocal disease. Mutations in normal samples had low allele frequencies compared to tumor mutations (p<2.2*10^−16^). Furthermore, significant differences in the type of nucleotide changes between tumor, normal and shared mutations (*p*=2.7*10^−8^) were observed, and mutations in APOBEC context were observed primarily among tumor mutations (*p*=0.026). No differences in functional impact between normal, shared and tumor mutations were observed (*p*=0.23). Overall, these findings support the theory of multiple fields in the bladder, and document non-tumor specific driver mutations to be present in normal appearing bladder tissue.

## Introduction

By applying whole exome sequencing and deep targeted sequencing on bladder tumors, it was recently shown that tumors developed years apart in the same patients share multiple mutations and hence are clonally related^1-3^. Furthermore, apparently normal urothelium has been documented to contain mutations with low allele frequencies (∼3%) that are typically observed at high frequencies in tumors (clonal mutations)^1-3^. Multiple studies have investigated genomic alterations in normal appearing bladder tissue from cystectomy specimens, however using technologies that do not allow detection of low-frequency mutations. The genomic alterations observed in these studies include copy number alterations of chromosome 5, 9, 13, 16, and 17 as well as mutations or loss of *RB1* and *TP53*^4-9^. These findings corroborates the suggestions of the presence of field cancerization in the bladder. Similar results have been reported in other tissue types, where studies have revealed the presence of mutations in well-characterized cancer driver genes in apparently healthy tissue and pre-cancer lesions^10-13^.

Bladder cancer (BC) is multifocal in almost half of the cases with primary tumour and in more than 50% of the patients with recurrent non-muscle invasive BC (NMIBC)^14^. Moreover, recurrent BC is common as the majority of the patients with non-muscle invasive BC (NMIBC) relapse within five years^15,16^. Approximately 75% of patients with BC present with NMIBC, and 5-25% of these will progress to muscle-invasive bladder cancer (MIBC)^16,17^. Multifocality and the frequent recurrences of BC are hypothesized to originate from field cancerization of the bladder urothelium^18^. This concept was first described in oral squamous epithelium in 1953 by Slaughter et al. as an explanation of the high local recurrence rate of oral cancers^19^. More recent, field cancerization has been described as an underlying mechanism for tumor development in various cancer types, including BC^20^.

Field cancerization is understood as one or more areas, or fields, with mutated cells. Normal cell lineages acquire mutations that are positively selected for in the microenvironment of an otherwise healthy organ. Consequently, the mutant clone can grow to produce fields of a monoclonal origin that predispose to malignant growth within these transformed areas. The transformed cells may appear normal or dysplastic^20,21^. Thomsen et al proposed a theory of multiple fields being present in the bladder^2^ where parallel expansion of different mutated stem cells might lead to multiple transformed fields intermixed in the bladder urothelium. Tumors will mirror the genetic alterations from the field from which it arose. This theory may explain the low frequencies of mutations observed in normal samples^2^.

In our previous study of bladder cancer field cancerization we analyzed mutations in adjacent normal tissue restricted to mutations observed in the tumor samples, and consequently, non-tumor specific mutations were not investigated^2^. In this study, we characterized mutations in normal appearing urothelium adjacent to tumors by deep targeted sequencing. We detected high-impact mutations in known driver genes that were not observed in the tumor. Furthermore, we observed mutations shared between tumor and normal samples (tumor field effect) as well as mutations specific to the tumors (mutations acquired later in development).

## Results

We performed deep-targeted sequencing of DNA obtained from four patients (patients 1 to 4) with advanced bladder cancer, treated with radical cystectomy (see Supplementary Fig. S2 and Supplementary Table S3 for detailed disease courses). From each patient, DNA was procured from bulk tumor biopsies (n=2-7) and laser microdissected (LMD) biopsies of normal appearing urothelium (n=6-11) (See Supplementary Table S1 for overview of samples and sequencing information). Individual bulk tumor samples were previously analyzed by whole exome sequencing (WES) followed by deep targeted amplicon sequencing of LMD tumor and normal samples guided by the original WES of bulk tumor^2^. In this present study, we expand on our previous study to include the analysis of mutations uniquely present in normal appearing adjacent tissue by deep targeted sequencing (Figure 1a).

**Figure 1:**
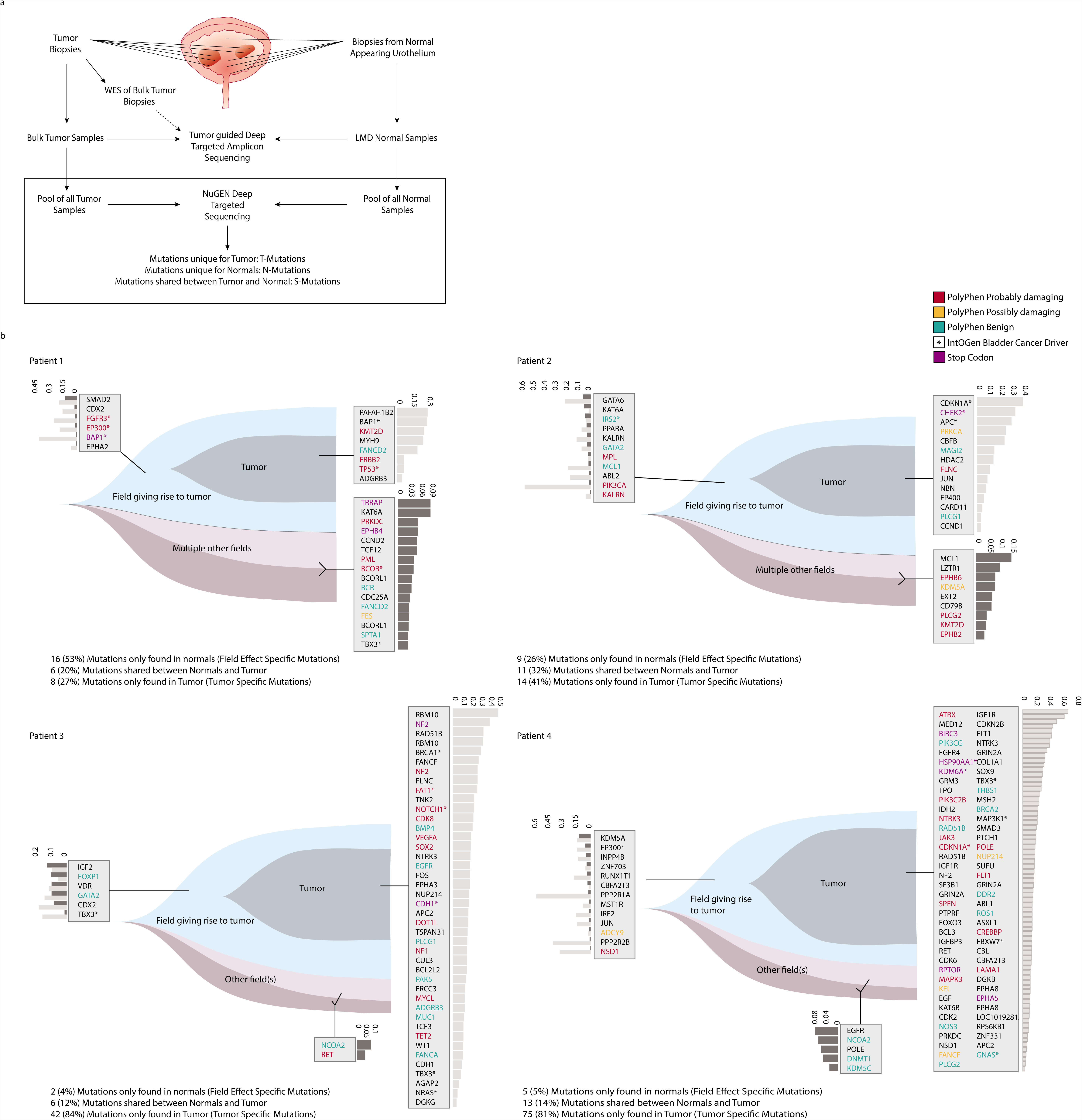
Analysis of field cancerization in four patients. (a) Study design. Upper part: analyses performed previously. WES was performed on bulk tumor samples. Multiple tumor and normal biopsies were laser microdissected (LMD) and subjected to deep targeted amplicon sequencing guided by the bulk tumor WES. Lower part: present study (black box). Tumor and normal DNA samples were pooled and subjected to deep targeted amplicon sequencing. Mutation calls were analyzed and grouped into T-Mutations, N-Mutations, and S-Mutations. (b) Analysis of patients 1-4. Field cancerization visualized using T-Mutations, N-Mutations, and S-Mutations. Gene names and allele frequencies (AF) are displayed. AFs are illustrated as light grey bars (AF measured in tumor) and dark grey (AF measured in normal).

### Deep targeted sequencing

Extracted DNA from tumors and LMD normal samples was pooled resulting in one pool of tumor DNA (tumor pool) and one pool of normal DNA (normal pool) from each of the four patients. We performed deep targeted amplicon sequencing of 509 cancer genes on both pools and on matched leukocyte DNA as reference. We obtained an average target read depth of 634x (range: 360-1073). Following sequence read consolidation (UID error correction)^22^ the average target read depth was 69x (range: 36-129). In total, after filtering, we identified 30-93 mutations in the samples from the four patients. Of these, 2-16 were unique for pools of normal samples (N-Mutations), 8-75 were unique for tumor pools (T-Mutations), and 6-13 were shared between tumor pools and normal pools (S-Mutations)(Figure 1b).

### Analysis of field cancerization

Patients 1 and 2 presented with multifocal disease, whereas patients 3 and 4 had unifocal disease. In patients 1 and 2, 39% (25/64) of the mutations were N-Mutations, and 34% (22/64) were S-Mutations. Mutations called in patients 3 and 4 were mainly T-Mutations, with only 5% (7/143) being N-Mutations and 13% (19/143) S-Mutations - indicating that uni- and multifocal patients may show different levels of field cancerization. Mutations in known BC driver genes were detected in both N-, S- and T-Mutation groups, most of them being among T-Mutations. However, in patient 1, two N-mutations were observed in bladder cancer driver genes. Damaging mutations were present in all N-, S- and T-Mutation groups. We detected the introduction of premature stop codons, mainly in the T-Mutation group. However, for patient 1 premature stop codons were solely observed within the N- and S-Mutations. Mutation allele frequencies (AFs) varied for the different mutations detected but were generally low for N-Mutations and high for T-Mutations. See Figure 1b and **Table 1** for details.

**Table 1:**
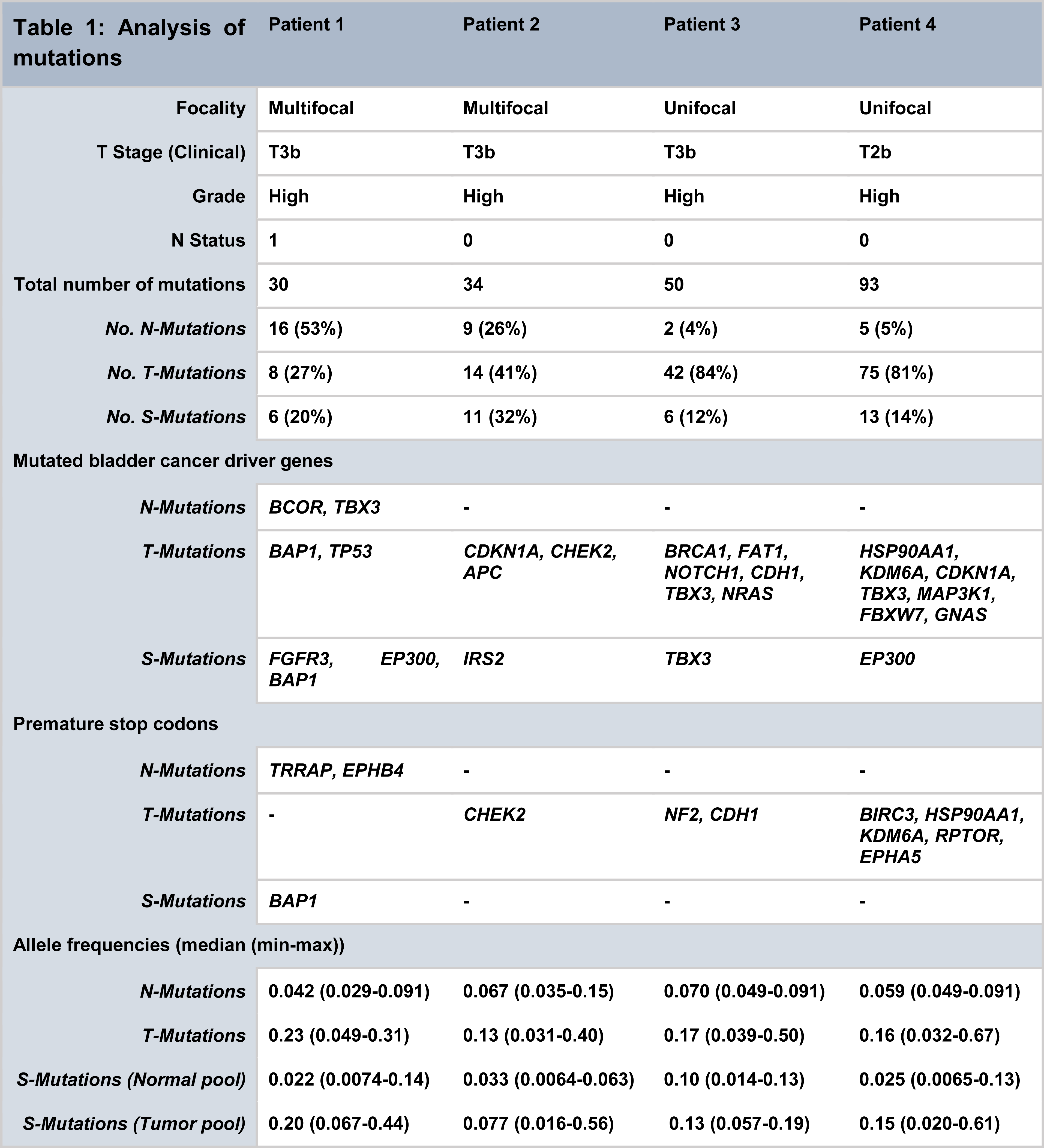

Interestingly, we observed N-Mutations in genes known to have a role in cancer development. To corroborate our findings, we investigated the genes affected by non-synonymous mutations in 1889 patients with a total of 1934 samples from 11 different BC studies using cBioPortal. In total, 0.6% to 23% (mean 4%) of the bladder tumors harbored mutations in the same set of genes. The six most frequently non-synonymous mutated N-Mutation genes in the BC datasets were KMT2D (23%), SPTA1 (8%), TRRAP (7%), PRKDC (6%), POLE (4%), and KDM5A (4%).

### Validation of mutations by WES and ddPCR

Validation of mutations was performed in a two-step process. Firstly, WES data of tumor samples was used to validate mutations detected by our deep targeted sequencing approach. In general, we observed consistency in AFs measured by the two platforms, and most positions were covered across all samples (Spearman correlation=0.77, *p-val*=2.2*10^−16^) (Figure 2a and 2b).

**Figure 2:**
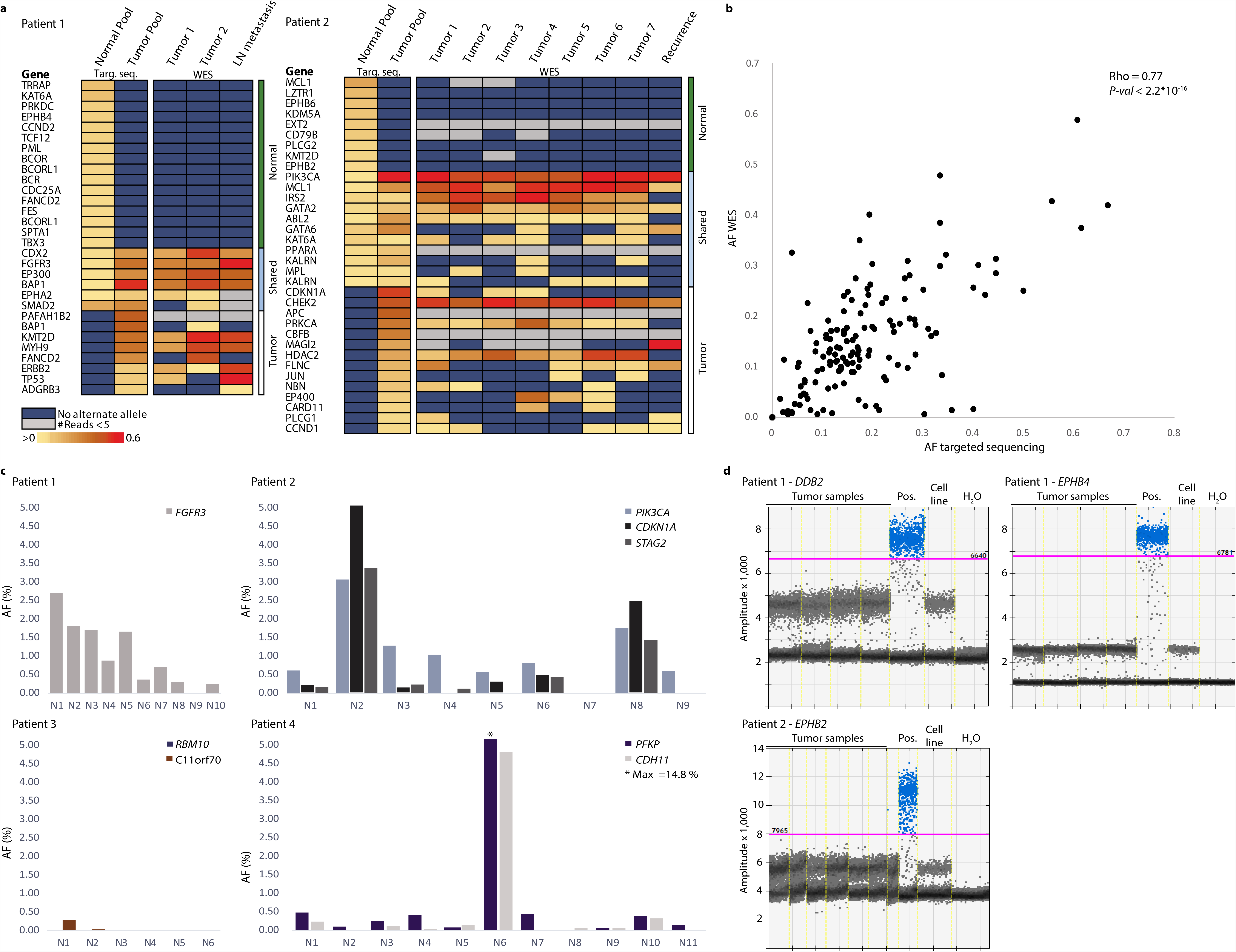
Validation of mutations. (**a**) All mutations were evaluated in previously generated WES data from tumors, recurrences, and metastases from the four patients (patients 1 and 2 shown, patients 3 and 4 in Supplementary Fig. S3). Obtained AFs are marked (yellow to red ranging from >0 to 0.6). For WES data, a minimum of five reads at a given position were required for validation (indicated in grey). Dark blue indicates no alternate alleles on the position. LN = lymph node. Targ. seq. = Targeted sequencing.(**b**) AFs obtained by cancer panel sequencing of tumor compared to mean AFs from WES on tumor samples from all four patients. Recurrences and metastases were excluded from calculation of the mean as these samples were not included in the tumor pools. Spearman correlation was calculated. (**c**) Validation of previously identified tumor mutations^2^ by ddPCR on DNA from normal samples. Multiple assays for specific mutations were included for the four patients and the fraction of mutated sequences identified using ddPCR is shown (%). * indicates that the value is out of scale (max value = 14.8%). (**d**) Validation of absence of N-Mutations in DNA from tumor samples by ddPCR analysis. A positive control (synthesized oligo) for each assay was included as well as negative controls (H2O and HT1197 bladder cancer cell line). The purple line indicates cutoff set for positive droplets. Droplets positive for mutation are marked in blue and negative droplets are indicated by grey.

Secondly, we used ddPCR to validate the presence/absence of selected alterations in normal and tumor samples. Eight mutations previously observed in tumor and normal samples^2^ and three additional N-Mutations were chosen for ddPCR validation. For every patient, tumor mutations were analyzed by ddPCR in 6-11 samples from the normal appearing urothelium. Except for a deletion in *RBM10*, the tumor alterations were detected at low frequencies in normal samples (Figure 2c). AFs from ddPCR were compared to deep targeted amplicon sequencing of the same samples and a correlation coefficient of 0.93 was observed. For N-Mutation analysis, DNA extracted from 4-7 tumor areas were analyzed and none of the mutations were detected in any of the tumor samples analyzed by ddPCR (Figure 2d), which validated the normal tissue specificity

### Analysis of mutational context

We performed a combined analysis of the mutations detected in the four patients as the individual patients harbored too few mutations for robust statistical analyses. We observed a significant difference in the six single-base substitutions between the three groups of mutations (*p*=2.7*10^−8^, Fisher’s Exact Test): 58% of T-Mutations were C>T changes compared to 40% of both N- and S-Mutations. Furthermore, we observed no T>G mutations in N-Mutations, whereas 40% of S-Mutations and 1.5% of T-Mutations were T>G base pair substitutions. C>G mutations were present among N-Mutations and T-Mutations at 25% and 22% frequency, respectively, compared to 3% in S-Mutations (Figure 3a). C>T mutations have been associated with various signatures, including the age-dependent signature 1 and the APOBEC-related signature 2. C>G substitutions have been attributed to signature 13 (APOBEC related), which is commonly observed in BC^13,23-25^.

**Figure 3:**
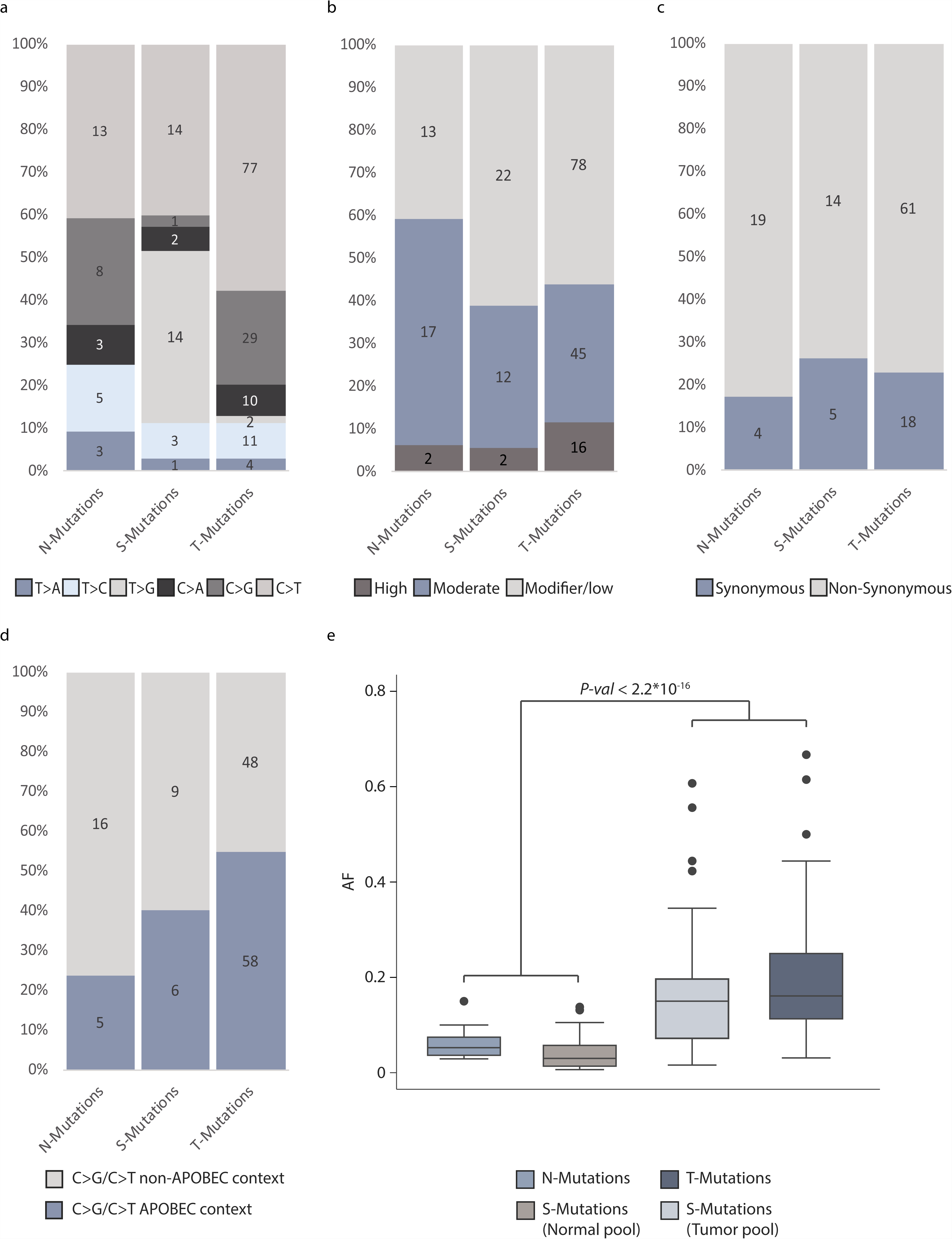
Analysis of mutational context, impact and frequency. All analyses were performed on the combined set of mutations from all patients. The total number of mutations in each category is indicated. (**a**) The six single-base substitutions counted among N-, T-, and S-Mutations. (**b**) Predicted impact of mutations among N-, T-, and S-Mutations grouped into high, moderate, low/modifier impact. (**c**) Predicted impact of mutations in N-, T-, and S-Mutations grouped into synonymous and non-synonymous (mutations predicted to have a high or moderate impact) mutations (**d**) Number of C>G and C>T mutations among N-, T-, and S-Mutations in APOBEC context. (**e**) Allele frequencies from N-, T-, and S-Mutations. For S-Mutations, allele frequencies are measured both in the normal samples and in the tumor samples and both are indicated.

We observed no difference in the functional impact of the mutations observed in the three mutation categories. This was observed both when assessing mutations categorized as being of high, moderate, or low/modifier impact by the SNPEff software (*p*=0.23, Fisher’s Exact Test), and when analyzing synonymous and non-synonymous mutations (*p*=0.77, Fisher’s Exact Test) (Figure 3b and 3c).

Next, we assessed the proportion of APOBEC related mutagenesis. C>T/G mutations in a TCW context, where W is either T or A, were evaluated as representing the APOBEC signature^26^. We observed a significant difference between the proportion of N-, S-, and T-Mutations in APOBEC related context (*p*=0.0011, Fisher’s Exact Test). In addition, we observed a significant difference when comparing C>T/C>G mutations in an APOBEC-related context and C>T/C>G in non-APOBEC related context in N-, S-, and T-Mutations (*p*=0.026, Fisher’s Exact Test) (Figure 3d).

Finally, AFs for mutations in normal samples were significantly lower than for mutations in tumor samples (p<2.2*10^−16^, Unpaired T-test)(Figure 3d). There was no significant difference between AFs for T-Mutations and S-Mutations measured in the tumor pool (*p*=0.09, Unpaired T-test).

## Discussion

Here we characterized the field cancerization in four patients with advanced BC and addressed the question of multiple mutated fields being present within the bladder. Field cancerization was observed in all four patients analyzed, being more pronounced in patients with multifocal disease compared to patients with unifocal disease. We found that the normal appearing urothelium harbored private mutations not detected in the tumor samples. We suggest that these mutations represent one or more fields that have not lead to tumor development. Additionally, we detected mutations that were shared between normal and tumor samples, representing mutations from the field developing into a tumor. Mutations unique for tumor samples were also present, indicating further genomic evolution of the tumor after initial development from the field.

Different origins of these mutated cells have been proposed^21^. These include intraepithelial migration and/or luminal seeding of carcinoma cells from existing tumors followed by implantation of the carcinoma cells – eventually giving rise to recurrent tumors. Another theory is that the field develops before the tumor from an altered stem cell embedded in the urothelium. Following this, the altered clone can expand, leading to a population of mutated daughter cells forming a cancerized field^20,21^.

Our analysis showed that mutations were present at low frequencies in the normal appearing samples. This could be explained by the seeding of tumor cells from existing tumors, resulting in the presence of a few mutated tumor cells in normal samples. Also, it could be due to some tumor cells migrating through the epithelial layer^18^. However, these explanations do not explain the presence of mutations unique for the normal samples. Therefore, another possible explanation for the presence of low frequency mutations in normal samples is that a few mutated cells are intermixed either with normal cells or with other differently transformed cells. Different mutated cell lineages can arise if more self renewing cells (e.g. stem cells) are mutated in different ways and expand in parallel, creating multiple transformed fields^2,27,28^. This theory may explain the presence of normal specific mutations. If recurrent tumors develop from fields that arose from the same mutated stem cell, these will be clonally related^2^. This could hence explain the clonal origin of metachronous bladder tumors^1^ as well as paired upper tract and bladder urothelial tumors^29^.

Two studies from Martincorena et al.^11,13^ have revealed the presence of non-tumor specific mutations in normal tissue from esophagus and skin, respectively. These results indicate that the field arise prior to eventual tumor development and that normal cells harbor mutations without necessarily developing into a tumor. To our knowledge, no previously published studies have focused on mutations in normal appearing bladder tissue without being restricted to mutations observed in the tumor. Our study was performed on normal appearing bladder tissue for non-tumor guided detection of mutations. In order to detect these low-frequency mutations in normal samples, it is necessary to perform deep sequencing. Furthermore, to differentiate low frequency mutations from common sequencing errors, error correction methods, such as the inclusion of UIDs^22^, should be included in the sequencing and subsequent analyses.

We observed that the expected impact of N-, S- and T-Mutations was the same across all three groups. We would expect S-Mutations and T-Mutations to have a higher impact than N-Mutations, as these two groups drive initial tumor formation and later tumor evolution. In the Martincorena et al studies, high impact mutations, missense mutations, and cancer driver mutations were observed in normal tissue from non-cancerous individuals^11,13^. Consequently, these findings may imply that tumor formation is more dependent on the affected genes, combination of genes, and the order in which mutations occur^30^. Additionally, from our analysis it is not possible to know how many mutations are present in the individual cell, and future studies utilizing single cell sequencing are needed to delineate the genomic changes per cell.

In addition, we observed that mutations in APOBEC context were mainly present in the T-Mutation group. This is in concordance with other studies that have suggested that APOBEC mediated mutagenesis is a late event in tumor evolution^31,32^. Furthermore, most of the non-APOBEC related C>T mutations observed in the normal samples were found in a CpG context (7/11) and may hence be related to the age related signature 1, in accordance with the fact that mutations accumulate in normal cells over time^24^.

We hypothesize that field cancerization may have prognostic and predictive value. However, as stated previously, results from our and other studies have shown that mutations do indeed occur in normal cells without leading to cancer development. This may affect screening initiatives for early detection of cancer using e.g. analysis of mutated DNA in urine and plasma. Detection of high impact mutations might not imply that patients have cancer. A recent study detected mutations in cfDNA from individuals without cancer, documenting the need for using tumor guided approaches^33^.

In conclusion, this study sheds light on the field cancerization in BC, and documents that non-tumor specific mutations are present in normal appearing tissue. It will be necessary to analyze tissue from additional patients to be able to better describe the field cancerization and its role in tumor development, disease recurrence and aggressiveness, and e.g. BCG treatment efficacy. Moreover, novel methods for single cell analysis may be powerful supplements to better understand the biology of field cancerization.

## Patients and methods

### Clinical samples

Patients included in the study were diagnosed with primary BC and underwent open radical cystectomy and extended lymph node dissection to the aortic bifurcation. The patients had not received neoadjuvant chemotherapy or radiation therapy before cystectomy. Tissue biopsies were embedded in TissueTek OCT^TM^ Compound (Sakura, Finetek, Vaerloese, Denmark), snap-frozen in liquid nitrogen and stored at −80 °C. Two to seven biopsies were obtained from tumors from each patient together with six to 12 biopsies taken throughout the normal appearing urothelium. Blood samples were stored in EDTA tubes at −80 °C. Areas of tumor and normal urothelium were LMD for all patients to ensure cell content specificity of the samples. LMD and DNA extraction from bulk and LMD samples and blood samples were performed as described previously^2^. Patients were treated at Aarhus University Hospital in 2014 and provided informed written consent. The study was approved by The Danish National Committees on Health Research Ethics (#1300174). All methods in the study were carried out in accordance with the approved guidelines and regulations.

### Targeted sequencing and data processing

Targeted sequencing was performed on pools of normal samples and pools of tumor samples using the NuGEN Ovation^®^ Cancer Panel 2.0 Target Enrichment System (509 genes; NuGEN Technologies)^34^. DNA from normal samples and tumor samples from each patient was pooled prior to library generation in order to obtain enough input material. Tumor pools for all patients consisted of 1:1 amounts of bulk tumor DNA. Libraries were prepared from 500 ng DNA (Qubit), as previously described^22^. Libraries were amplified using 21 PCR cycles and subsequently pooled eight at a time and single-end sequenced (150bp) on an Illumina NextSeq 500 (High output).

Sequencing data was aligned and mapped, as previously described^22^. In brief, reads with identical UIDs and mapping positions were collapsed to create high confidence consensus reads. If less than three reads shared UIDs and mapping positions, they were discarded. Mutations were called using MuTect2. Mutations identified in pools of normal samples and/or pools of tumor samples were assessed using bam-readcount in previously generated WES data. WES data was obtained from tumor and leukocyte samples from the same patients and processed as previously described^1,2^. Moreover, mutations identified in pools of normal samples were assessed in the associated pools of tumor samples and vice versa.

### Filtering of mutations

Initially, mutations were categorized in three different sets based on whether they were called (MuTect2) or observed (pileup tools) only in normal samples (Normal specific mutations - N-Mutations), only in tumor samples (Tumor specific mutations or T-Mutations) or in both pools (Shared mutations or S-Mutations) using the cancer panel sequencing (Supplementary Fig. S1). To ensure normal sample specificity, initial N-Mutations were evaluated in previously generated WES data. Mutations were discarded if present with two or more alternate reads in any of the corresponding tumor samples.

Any positions with more than two alleles were excluded and all remaining mutations were reviewed manually using the Integrative Genomics Viewer (IGV)^35^.

### Functional assignment

We identified mutations in known BC driver genes defined in IntOGen (BBGLab)^36^ and assigned the functional impact to mutations using PolyPhen-2 and snpEff v4.3^37,38^.

### Digital Droplet PCR (ddPCR)

For the validation of N-Mutations, an oligo covering the whole mutated amplicon of interest (positive control) was designed due to insufficient sample amounts. ddPCR and data analysis were performed as previously described^39^. Assays targeting regions on chromosome 16 and 3 were used for quantification of total DNA copies as these regions are rarely subject to copy number alterations in BC^37^. Primer and probe sequences are listed in Supplementary Table S2.

### Statistical analysis

The Shapiro-Wilk test or Quantile-Quantile plot (QQ-plot) was used to test for normality of the data. Statistical analyses were performed using unpaired t-test on log-transformed parametric data with Welch correction for data with significantly different standard deviations. For categorical variables, Fisher’s Exact test was used. Correlation was calculated using Spearman. Statistical significance was set at *p<*0.05. All statistical analyses were performed using R (R version 3.5.1).

### Data availability

The raw sequencing datasets generated during the current study are not publicly available due to local Danish legislation on data sharing. However, processed datasets are available from the corresponding author on reasonable request.

## Acknowledgements

This work was funded by the Danish Cancer Research Foundation, the Danish Cancer Society, and the Danish Cancer Biobank (grant numbers are not applicable).

## Contributions

M.B.H.T. and J.B.J. collected patient material. T.S. performed clinical follow up. M.B.H.T. performed LMD experiments and DNA-extraction. T.S. and I.N. performed experimental work. T.S., E.C., and P.L. performed bioinformatic analyses. L.D., I.N., P.L., and T.S. designed the study and interpreted data. T.S. drafted the manuscript with input from all authors.

## References

1. Lamy, P. et al. Paired Exome Analysis Reveals Clonal Evolution and Potential Therapeutic Targets in Urothelial Carcinoma. Cancer Res. 76, 5894–5906 (2016).

2. Thomsen, M. B. H. et al. Comprehensive multiregional analysis of molecular heterogeneity in bladder cancer. Sci. Rep. 7, 11702 (2017).

3. Nordentoft, I. et al. Mutational context and diverse clonal development in early and late bladder cancer. Cell Rep. 7, 1649–1663 (2014).

4. Chaturvedi, V. et al. Superimposed histologic and genetic mapping of chromosome 17 alterations in human urinary bladder neoplasia. Oncogene 14, 2059–2070 (1997).

5. Czerniak, B. et al. Superimposed histologic and genetic mapping of chromosome 9 in progression of human urinary bladder neoplasia: implications for a genetic model of multistep urothelial carcinogenesis and early detection of urinary bladder cancer. Oncogene 18, 1185–1196 (1999).

6. Czerniak, B. et al. Genetic modeling of human urinary bladder carcinogenesis. Genes Chromosomes Cancer 27, 392–402 (2000).

7. Kram, A. et al. Mapping and genome sequence analysis of chromosome 5 regions involved in bladder cancer progression. Lab. Invest. 81, 1039–1048 (2001).

8. Majewski, T. et al. Understanding the development of human bladder cancer by using a whole-organ genomic mapping strategy. Lab. Invest. 88, 694–721 (2008).

9. Yoon, D. S. et al. Genetic mapping and DNA sequence-based analysis of deleted regions on chromosome 16 involved in progression of bladder cancer from occult preneoplastic conditions to invasive disease. Oncogene 20, 5005–5014 (2001).

10. Weaver, J. M. J. et al. Ordering of mutations in preinvasive disease stages of esophageal carcinogenesis. Nat. Genet. 46, 837–843 (2014).

11. Martincorena, I. et al. Tumor evolution. High burden and pervasive positive selection of somatic mutations in normal human skin. Science 348, 880–886 (2015).

12. Wood, H. M. et al. The clonal relationships between pre-cancer and cancer revealed by ultra-deep sequencing. J. Pathol. 237, 296–306 (2015).

13. Martincorena, I. et al. Somatic mutant clones colonize the human esophagus with age. Science (2018). doi:10.1126/science.aau3879

14. Ströck, V. & Holmäng, S. A Prospective Study of the Size, Number and Histopathology of New and Recurrent Bladder Tumors. Urology Practice 2, 260–264 (2015).

15. Sylvester, R. J. et al. Systematic Review and Individual Patient Data Meta-analysis of Randomized Trials Comparing a Single Immediate Instillation of Chemotherapy After Transurethral Resection with Transurethral Resection Alone in Patients with Stage pTa-pT1 Urothelial Carcinoma of the Bladder: Which Patients Benefit from the Instillation? Eur. Urol. 69, 231–244 (2016).

16. Kamat, A. M., Bagcioglu, M. & Huri, E. What is new in non-muscle-invasive bladder cancer in 2016? Turk J Urol 43, 9–13 (2017).

17. Babjuk, M. et al. EAU Guidelines on Non–Muscle-invasive Urothelial Carcinoma of the Bladder: Update 2016. Eur. Urol. 71, 447–461 (2017).

18. Höglund, M. Bladder cancer, a two phased disease? Semin. Cancer Biol. 17, 225–232 (2007).

19. Slaughter, D. P., Southwick, H. W. & Smejkal, W. ‘Field cancerization’ in oral stratified squamous epithelium. Clinical implications of multicentric origin. Cancer 6, 963–968 (1953).

20. Curtius, K., Wright, N. A. & Graham, T. A. An evolutionary perspective on field cancerization. Nat. Rev. Cancer 18, 19–32 (2018).

21. Braakhuis, B. J. M., Tabor, M. P., Kummer, J. A., Leemans, C. R. & Brakenhoff, R. H. A genetic explanation of Slaughter’s concept of field cancerization: evidence and clinical implications. Cancer Res. 63, 1727–1730 (2003).

22. Christensen, E. et al. Optimized targeted sequencing of cell-free plasma DNA from bladder cancer patients. Sci. Rep. 8, 1917 (2018).

23. Alexandrov, L. B. et al. Clock-like mutational processes in human somatic cells. Nat. Genet. 47, 1402–1407 (2015).

24. Alexandrov, L. B. et al. Signatures of mutational processes in human cancer. Nature 500, 415–421 (2013).

25. Ju, Y. S. The mutational signatures and molecular alterations of bladder cancer. Transl. Cancer Res. 6, S689–S701 (2017).

26. Roberts, S. A. et al. An APOBEC cytidine deaminase mutagenesis pattern is widespread in human cancers. Nat. Genet. 45, 970–976 (2013).

27. Höglund, M. On the origin of syn- and metachronous urothelial carcinomas. Eur. Urol. 51, 1185–93; discussion 1193 (2007).

28. Pardal, R., Clarke, M. F. & Morrison, S. J. Applying the principles of stem-cell biology to cancer. Nat. Rev. Cancer 3, 895–902 (2003).

29. Audenet, F. et al. Clonal Relatedness and Mutational Differences between Upper Tract and Bladder Urothelial Carcinoma. Clin. Cancer Res. (2018). doi:10.1158/1078-0432.CCR-18-2039

30. Levine, A. J., Jenkins, N. A. & Copeland, N. G. The Roles of Initiating Truncal Mutations in Human Cancers: The Order of Mutations and Tumor Cell Type Matters. Cancer Cell 35, 10–15 (2019).

31. Jamal-Hanjani, M. et al. Tracking the Evolution of Non-Small-Cell Lung Cancer. N. Engl. J. Med. 376, 2109–2121 (2017).

32. Jolly, C. & Van Loo, P. Timing somatic events in the evolution of cancer. Genome Biol. 19, 95 (2018).

33. Phallen, J. et al. Direct detection of early-stage cancers using circulating tumor DNA. Sci. Transl. Med. 9, (2017).

34. Li, H. & Durbin, R. Fast and accurate short read alignment with Burrows-Wheeler transform. Bioinformatics 25, 1754–1760 (2009).

35. Robinson, J. T., Thorvaldsdóttir, H., Wenger, A. M., Zehir, A. & Mesirov, J. P. Variant Review with the Integrative Genomics Viewer. Cancer Res. 77, e31–e34 (2017).

36. Gonzalez-Perez, A. et al. IntOGen-mutations identifies cancer drivers across tumor types. Nat. Methods 10, 1081–1082 (2013).

37. Schepeler, T. et al. A high resolution genomic portrait of bladder cancer: correlation between genomic aberrations and the DNA damage response. Oncogene 32, 3577–3586 (2013).

38. Cingolani, P. et al. A program for annotating and predicting the effects of single nucleotide polymorphisms, SnpEff: SNPs in the genome of Drosophila melanogaster strain w1118; iso-2; iso-3. Fly 6, 80–92 (2012).

39. Birkenkamp-Demtröder, K. et al. Monitoring Treatment Response and Metastatic Relapse in Advanced Bladder Cancer by Liquid Biopsy Analysis. Eur. Urol. 73, 535–540 (2018).

