## Supplementary Information for "Mutational analysis of field cancerization in bladder cancer"

**The following is included:**

Supplementary Fig. S1
Supplementary Fig. S2
Supplementary Fig. S3
Supplementary Table S1
Supplementary Table S2
Supplementary Table S3

**Supplementary Figures**

**Supplementary Fig. S1:** Filtering of mutations.

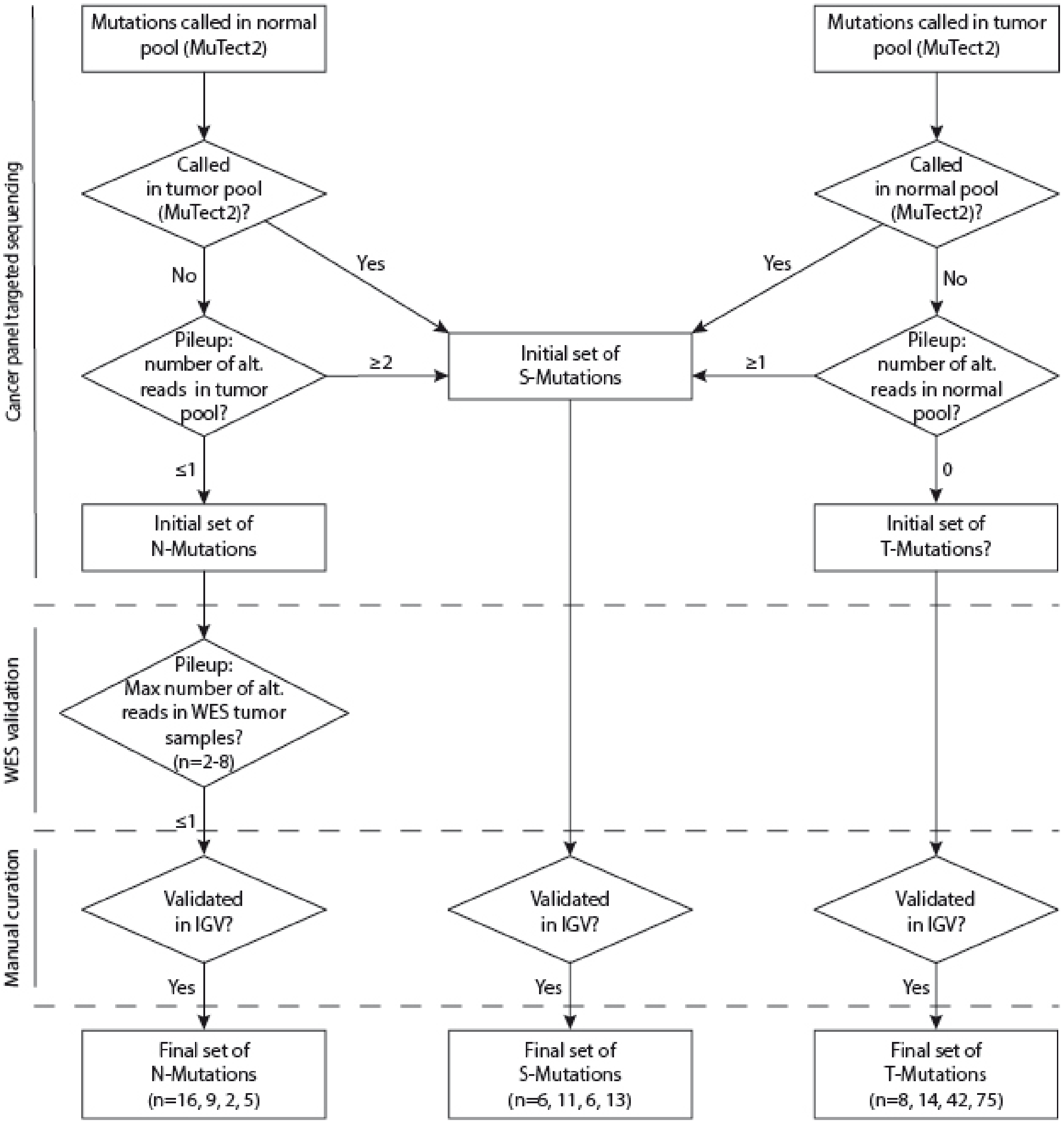


**Supplementary Fig. S1:** Filtering of mutation calls. Only mutations that pass the MuTect2 filter were considered for validation and analysis. Mutations were assessed in previously generated WES data and data obtained from the targeted sequencing of tumor and normal samples to ensure the presence and/or absence in other samples. Required number of reads for different groups is indicated. All mutations were evaluated manually using Integrative Genomics Viewer (IGV). Finally, mutations were grouped into N-Mutations, S-Mutations, and T-Mutations, N-Mutations being specific for normal samples, S-Mutations being shared between normal and tumor samples, and T-Mutations being exclusive for tumor samples.

**Supplementary Fig. S2:** Disease courses of the four patients.

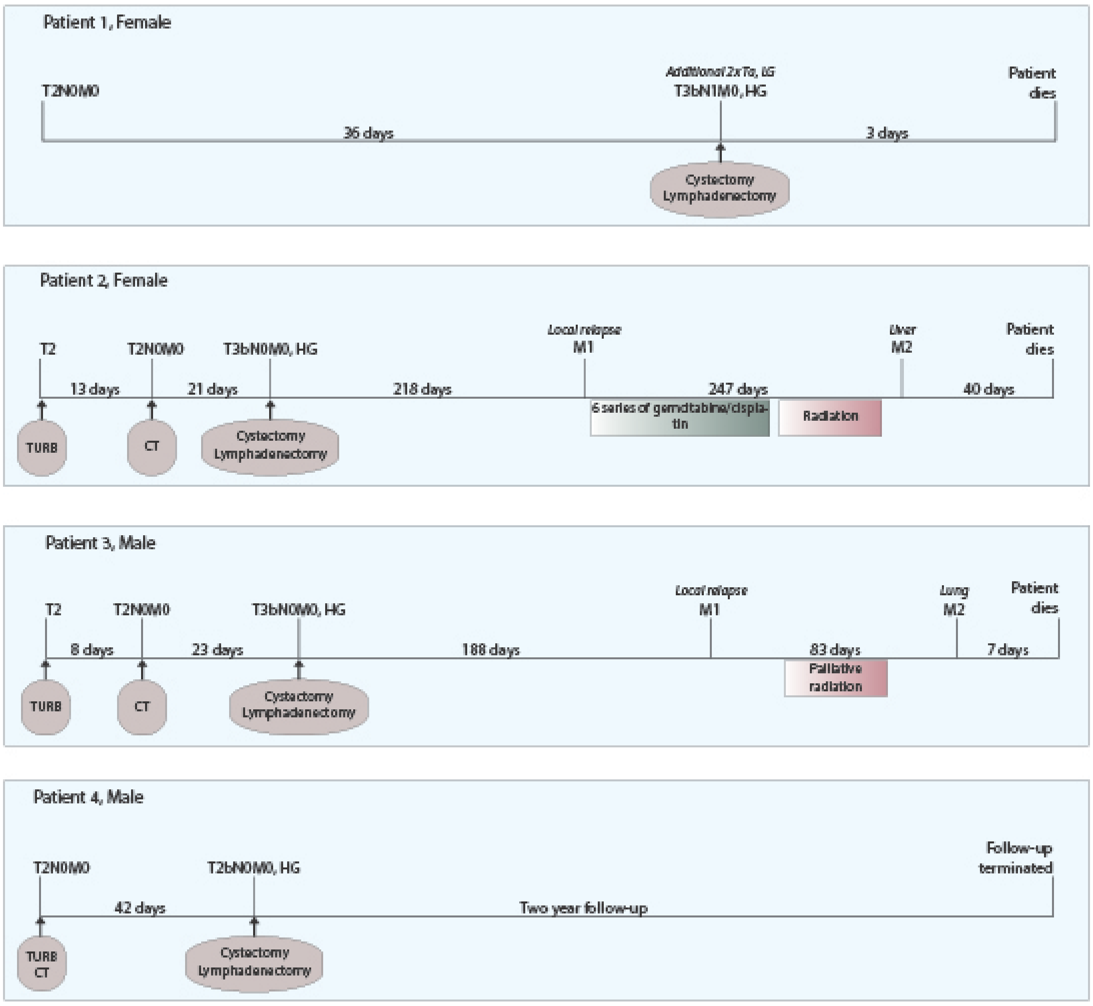


**Supplementary Fig. S2:** Disease courses of patients 1-4. TNM-stage for tumors at diagnosis as well as for tumors detected at follow-up visits and after cystectomy is indicated. Number of days between follow-up visits is annotated. Different treatments and examinations, including transurethral resection of bladder tumor (TURB), CT scans, cystectomy and lymphadenectomy, radiation, and chemotherapy are noted.

**Supplementary Fig. S3:** Comparison of allele frequencies obtained from the sequencing of normal and tumor samples using cancer panel targeted sequencing and from WES of tumor samples and recurrences from patients 3 and 4.

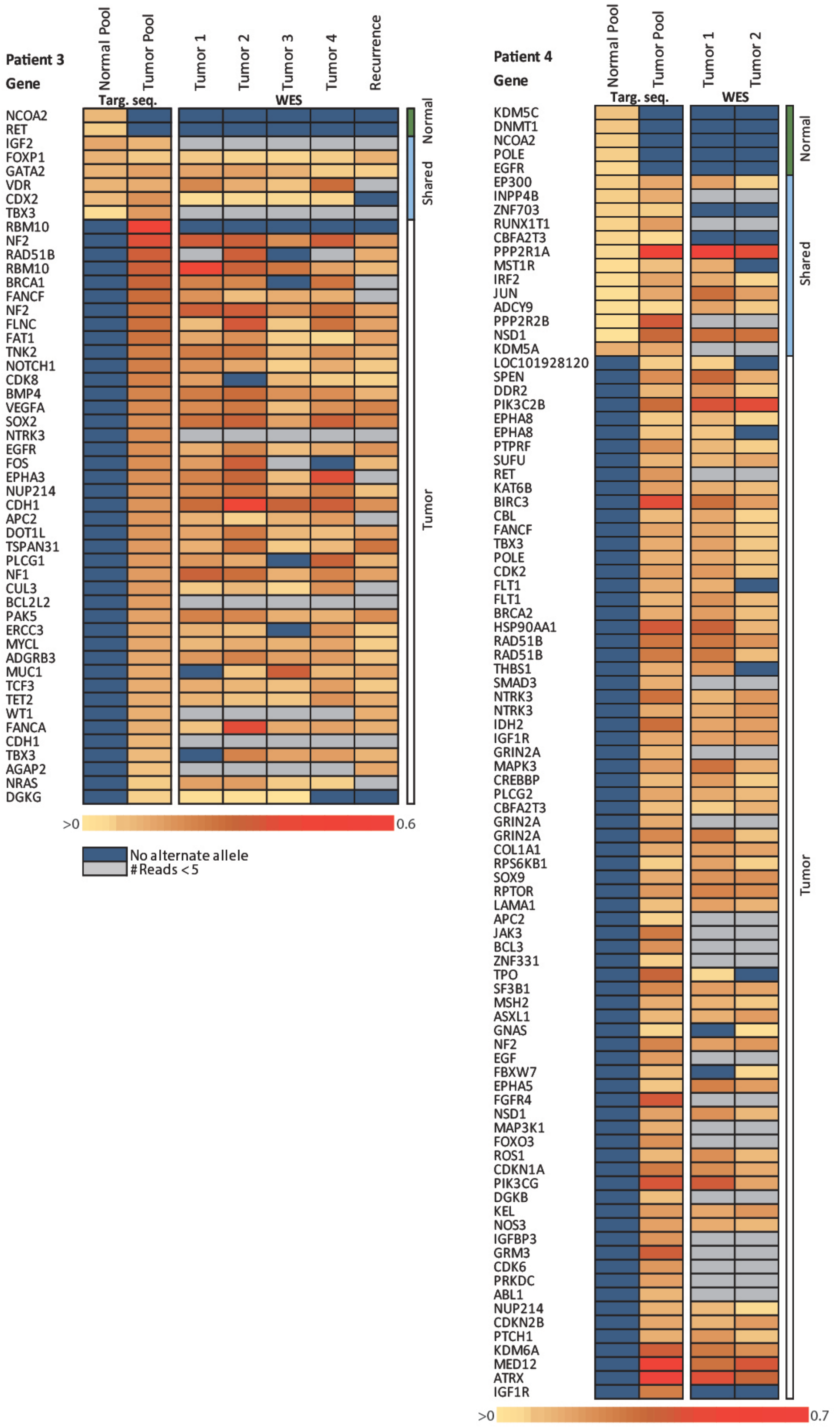

**Supplementary Fig. S3:** Comparison of different platforms. All mutations were evaluated in previously generated WES data from tumors and recurrences from patients 3 and 4. Obtained allele frequencies are marked (yellow to red ranging from >0 to 0.6/0.7). For WES data, a minimum of five reads were required and positions not fulfilling this indicated in grey. Dark blue indicates no alternate alleles on the position. Targ. seq. = Targeted sequencing.

### Supplementary Tables

**Supplementary Table S1:** Overview of samples, cancer panel targeted sequencing, and WES

| Samples | Patient 1 | Patient 2 | Patient 3 | Patient 4 |
|---|---|---|---|---|
| **Germline DNA** | Leukocytes | Leukocytes | Leukocytes | Leukocytes |
| **Samples in Normal Pool** | 10 | 9 | 6 | 11 |
| **Samples in Tumor Pool (T)** | 4 (2)[1] | 7 | 4 | 2 |
| **Lymph Node Metastasis (N)** | 1 | No | No | No |
| **Recurrence (M)** | No | Local | Local | No |
| **Type (T/N/M)** | FF/FFPE/-* | FF/-/FFPE | FF/-/FFPE | FF/-/- |

*FF = Fresh frozen tissue, FFPE = Formalin fixed paraffin embedded tissue

| Targeted Sequencing (NuGEN) | Patient 1 | Patient 2 | Patient 3 | Patient 4 |
|---|---|---|---|---|
| Number of Samples | 3 | 3 | 3 | 3 |
| Bait set | NuGEN Ovation® Cancer Panel 2.0 | NuGEN Ovation® Cancer Panel 2.0 | NuGEN Ovation® Cancer Panel 2.0 | NuGEN Ovation® Cancer Panel 2.0 |
| Mean Target Coverage (Normal Pool) (Raw/3UID) | 1065/79 (32-116)* | 1036/47 (18-70)* | 511/36 (15-53)* | 500/43 (18-63)* |
| Mean Target Coverage (Tumor Pool) (Raw/3UID) | 360/47 (19-67)* | 483/76 (27-104)* | 459/64 (25-93)* | 526/79 (27-110)* |
| Mean Target Coverage (Germline) (Raw/3UID) | 707/105 (49-152)* | 1073/129 (58-187)* | 442/52 (21-77)* | 445/70 (31-102)* |
| Total Mutations | 30 | 34 | 50 | 93 |
| Normal Mutations | 16 | 9 | 2 | 5 |
| Tumor Mutations | 8 | 14 | 42 | 75 |
| Shared Mutations | 6 | 11 | 6 | 13 |
| **WES (for mPileUp)** | **Patient 1** | **Patient 2** | **Patient 3** | **Patient 4** |
| Number of Exomes | 3 | 8 | 5 | 3 |
| Bait set | Nextera | SeqCap_EZ | Nextera | SeqCap_EZ |
| Mean Target Coverage (Tumor) | 86-107x | 31-82x | 52-113x | 62-65x |
| Mean Target Coverage (Germline) | 109x | 55x | 52x | 72x |

[1]For WES (previous study), two tumor samples were pooled into Tumor Pool 1, and two other tumor samples were pooled into Tumor Pool 2.
*Mean (lower quartile-upper quartile)

**Supplementary Table S2:** Primers and probes for ddPCR analyses

Primer sequences for detection of tumor mutations in normal samples by ddPCR

| Target Gene | Forward primer (5'-3') (Sigma Aldrich) | Reverse primer (5'-3') (Sigma Aldrich) | Dual-labelled probe (5'-3') (BioSearch) | Tm (ºC) |
|---|---|---|---|---|
| **FGFR3** | CTGGTGGAGGCTGACGAG | AAGAAGCCCACCCCGTAG | FAM-AGTGTGTGTGCAGGCATCCTCA-BHQ | 61.2 |
| **PIK3CA** | TGGAATGCCAGAACTACAATCTT | TCCAAAGCCTCTTGCTCAGT | FAM_TGACATTGCACACATTCGAAAGAC_BHQ1 | 59 |
| **CDKN1A** | GTCCGTCAGAACCCATGC | GCCATTAGCGCATCACAGT | FAM_TCTTCGGCCCCAGTGGAC_BHQ1 | 59 |
| **STAG2** | CAATTGTCATTAGGCTTAGCTTTTT | GGCATGTCTCCATTTTCGAT | FAM-CCCATGACCTTTGAAAGTGGGA_BHQ | 59 |
| **RBM10** | CTTCAGGCCTCCCAAGGT | CATAGCCCTCATCCTGTTGG | FAM-TCCGAGGAGCGCCGGTC_BHQ1 | 59 |
| **C11orf70** | AGCCCCCAGATGATTTCTTT | GGACTGAAAGGTCTGGTCAAA | FAM-CCCAGGTCCAGGCAGAATCAA_BHQ1 | 59 |
| **PFKP** | TTCTCCCCTAGGGCTACCAG | ACCTACCACTTGCAGGATGC | FAM-AGAGGCCGACTGGGAAAGTG_BHQ | 61.2 |
| **CDH11** | CCACCCCCTTCATCATCATA | GTGACCCTGAGAAGGCAAAA | FAM-TGATGTTCTCATGGACATCTTCTTCC_BHQ | 61.2 |
| **Chr16** | GGTACAAGTTGGTGTTCTGAG | GCCCAAGTGCAATCACAG | FAM-CTCTCTCAGTGGGTAGATTTGTTATCC-BHQ | - |
| **Chr16** | GGTACAAGTTGGTGTTCTGAG | GCCCAAGTGCAATCACAG | HEX-CTCTCTCAGTGGGTAGATTTGTTATCC-BHQ | - |
| **Chr3** | CTAGAAGATCTACCTCCAAGAGG | CCAGGCTGAAGCTATTCCAG | HEX-CTCATACATCTGGCATATGGGCTGG-BHQ | - |

Primers for validation of N-Mutations by ddPCR

| Target gene | Forward primer (5'-3')(Sigma Aldrich) | Reverse primer (5'-3')(Sigma Aldrich) | Dual-labelled probe (5'-3')(Sigma Aldrich) | Amplicon size | Tm (ºC) |
|---|---|---|---|---|---|
| **EPHB4** | GAAGCCACTCGCCCACAG | GTACTCAGCATTCCTCCCCAT | FAM-TGGTCCAGGAGAGGGTGTCAGGCCC_BHQ | 105 | 65 |
| **DDB2** | GCAGGTCCTAGCAGAAGATG | TTATGCTGGTGGAGGGTCC | FAM-AGATCCTGCCACAATGCCGCAGC_BHQ | 107 | 65 |
| **EPHB2** | ATTGTCATGTGGGAGGTGAT | GATCTTCCCTCTCCCATCTG | FAM-CCTACTCGGACATGACCAACCAGGATG_BHQ | 123 | 63 |

Sequences for positive controls

| Target gene | Forward primer (5’->3’) (Sigma Aldrich) | Amplicon size |
|---|---|---|
| **EPHB4** | GAAGCCACTCGCCCACAGAGCCAAAAGCTGAGTAGTGAGGCTGCCGCTGGTCCAGGAGAGGGTGT**C**AGGCCCTAGGGGGCAAGGATGGGGAGGAATGCTGAGTAC | 105 |
| **DDB2** | GCAGGTCCTAGCAGAAGATGTGACTCAGACTGCCTCTGGGTGGGGCTGGCTGGCCCACAGATCCTGCCAC**A**ATGCCGCAGCATCGTCAGGACCCTCCACCAGCATAA | 107 |
| **EPHB2** | TTGTCATGTGGGAGGTGATGTCCTATGGGGAGCGGCCCTACT**C**GGACATGACCAACCAGGATGTAAGTCTCCAAGGGGATAGGCAAGGCCTCTCTGGCCCACCAGATGGGAGAGGGAAGA | 120 |

**Supplementary Table S3:** Clinical and pathological information.

| | Patient 1 | Patient 2 | Patient 3 | Patient 4 |
|---|---|---|---|---|
| **Gender** | Female | Female | Male | Male |
| **Age at Diagnosis** | 70 | 69 | 73 | 86 |
| **T Stage (Clinical)** | 3b | 3b | 3b | 2b |
| **Grade** | High | High | High | High |
| **N Status** | 1 | 0 | 0 | 0 |
| **Focality** | Multifocal | Multifocal | Unifocal | Unifocal |
| **Treatment** | Cystectomy + Lymphadenoectomy | Cystectomy + Lymphadenoectomy | Cystectomy + Lymphadenoectomy | Cystectomy + Lymphadenoectomy |
| **Neo adj. Chemotherapy** | - | - | - | - |
| **Follow up (days)** | 4 | 506 | 279 | 731 |
| **Relapse** | - | Yes (Vagina) | Yes (Pelvis) | - |
| **Adj. Chemotherapy** | - | Gemcitabine/ Cisplatin | - | - |
| **Time to Relapse** | - | 218 | 142 | - |
| **Died of Disease** | Yes[a] | Yes | Yes | No |

[a]Patient died from complications following surgery
